# The p97 cofactor Ubxn7 facilitates replisome disassembly during S-phase

**DOI:** 10.1101/2021.12.16.472925

**Authors:** Zeynep Tarcan, Divyasree Poovathumkadavil, Aggeliki Skagia, Agnieszka Gambus

## Abstract

Complex cellular processes are driven by the regulated assembly and disassembly of large multi-protein complexes. In eukaryotic DNA replication, whilst we are beginning to understand the molecular mechanism for assembly of the replication machinery (replisome), we still know relatively little about the regulation of its disassembly at replication termination. Over recent years, the first elements of this process have emerged, revealing that the replicative helicase, at the heart of the replisome, is polyubiquitylated prior to unloading and that this unloading requires p97 segregase activity. Two different E3 ubiquitin ligases are now known to ubiquitylate the helicase under different conditions: Cul2^Lrr1^ and TRAIP. Here we have found two p97 cofactors, Ubxn7 and Faf1, which can interact with p97 during replisome disassembly in S-phase. Only Ubxn7 however facilitates efficient replisome disassembly through its interaction with both Cul2^Lrr1^ and p97. Our data therefore characterise Ubxn7 as the first substrate-specific p97 cofactor regulating replisome disassembly in vertebrates.

## INTRODUCTION

DNA replication is one of the most fundamental processes in life and its faultless execution is essential for normal cell fate. Until recently the final stage of eukaryotic DNA replication, the termination stage, was mostly unexplored. DNA replication initiates from thousands of replication origins; the positions within the genome where replicative helicases become activated and start unwinding DNA. These then move in opposite directions away from each other, creating two DNA replication forks. The replicative helicase is composed of Cdc45, Mcm2-7 hexamer and GINS complex (CMG complex) (1), and is positioned at the tip of replication forks forming a platform for replisome assembly (2). Once established, the replication forks replicate chromatin until they encounter forks coming in opposite directions from neighbouring origins. At this point the termination of replication forks takes place, with removal of the replisome from fully duplicated DNA being its final stage (3). In higher eukaryotes, (*X. laevis* egg extract, *C. elegans* embryos, mouse embryonic fibroblasts and human cells) replisome removal in S-phase is driven by the Cul2^LRR1^ ubiquitin ligase, which ubiquitylates Mcm7 within the terminated CMG helicase complex with lysine 48 (K48)-linked ubiquitin chains (4). The modified CMG is then recognized by the p97 segregase and removed from chromatin allowing for disassembly of the whole replisome built around the helicase (5). Any helicase complexes that fail to be unloaded in S-phase are alternatively unloaded in mitosis. Disassembly of these complexes in mitosis also depends on p97 segregase function, but this time requires the E3 ubiquitin ligase TRAIP (6). Consequently, disassembly in mitosis is driven by alternative species of ubiquitin chains that decorate Mcm7, namely K6- and K63-linked ubiquitin chains. TRAIP ubiquitin ligase can act also during S-phase: it interacts with the replisome and either ubiquitylates CMGs that converge at inter-strand crosslinks (ICLs) or ubiquitylates a protein crosslinked to DNA (DPC) that blocks progression of the replication fork (7,8).

p97 segregase (also referred to as VCP, Cdc48, CDC-48 and Ter94) is a hexameric AAA+ ATPase family member, that uses the free energy of ATP binding and hydrolysis to extract ubiquitylated protein targets from stable protein complexes, chromatin or membranes. As a result, p97 is essential for protein homeostasis in the cell and the dynamic behaviour of a multitude of multi-protein assemblies (9). The substrate specificity of p97 recognition is believed to be achieved by a number of regulatory cofactors (reviewed in (10)). In *C. elegans* embryos, the CDC-48 cofactors UFD-1/NPL-4 and UBXN-3 (Faf1 in higher eukaryotes) were shown to be required for replisome removal from chromatin in both S-phase and in mitosis (4,11). UFD-1/NPL-4 form a heterodimer, essential for most chromatin-related roles of p97, while UBXN-3 provides higher substrate specificity. Interestingly, Ufd1/Npl4 were also shown to interact with p97 and the replisome during replication termination in *Xenopus egg* extract (4). Despite this, additional cofactors of p97, which provide substrate specificity towards the terminated replisomes, are as yet to be determined in vertebrates.

Here, we have sought to identify p97 cofactors that are facilitating replisome disassembly during S-phase. While we were able to identify two new cofactors for this process, Ubxn7 and Faf1, our findings revealed that the Ubxn7 cofactor specifically, is crucial for efficient and fast disassembly of replisomes, as it creates bridges between the essential factors of this process: Cul2^Lrr1^, ubiquitylated Mcm7 and the p97 complexes.

## MATERIALS AND METHODS

### Inhibitors

MLN4924 (A01139, Active Biochem) was dissolved in DMSO at 20 mM and added to the extract 15 minutes after addition of sperm nuclei at 10 μM. NMS873 (17674, Cayman Chemical Company) was dissolved in DMSO at 10 mM and added to the extract 15 minutes after addition of sperm nuclei at 50 μM. Caffeine (C8960, Sigma) was dissolved in water at 100 mM and added to the extract along with demembranated sperm nuclei at 5 mM. Aphidicolin was dissolved in DMSO at 8 mM and added to the extract along with demembranated sperm nuclei at 40 μM.

### Recombinant proteins

Recombinant His-tagged ubiquitin and ubiquitin mutants were purchased from Boston Biochem, dissolved in LFB1/50 buffer (10% sucrose, 50 mM KCl, 40 mM HEPES pH 8.0, 20 mM K phosphate pH 8.0, 2 mM MgCl_2_,1 mM EGTA, 2 mM DTT, 1 μg/ml of each: aprotinin, leupeptin and pepstatin) at 10 mg/ml and used at 0.5 mg/ml *Xenopus laevis* egg extract.

pET28a-Ubxn7, pET28a-Ubxn7-P458G, pET28a-Ubxn7-L286E/A289Q/S293A vectors were used for protein expression in 2L of BL21-codon Plus (DE3)-RIPL (1 mM IPTG added at OD600 = 0.6, followed by incubation overnight at 20°C). Frozen bacterial pellets were lysed in LFB1/50 buffer (10% sucrose, 50 mM KCl, 40 mM HEPES pH 8.0, 20 mM K phosphate pH 8, 2 mM MgCl_2_, 1 mM EGTA, 2 mM DTT, 1 μg/ml of each: aprotinin, leupeptin and pepstatin) and supplemented with 1 mg/ml lysozyme and BitNuclease. After sonication, the lysate was clarified by centrifugation at 14,000 g for 30 min at 4°C, and supernatants containing soluble proteins were then incubated with 2 ml of pre-washed Super Ni-NTA Affinity Resin (SUPER-NINTA100, Generon) for 2 hours, rotating at 4°C. The beads were then washed 2x with 30 ml of LFB1/50 buffer and 2x with LFB1/200 (200 mM KCl), both supplemented with 20 mM imidazole. The beads were transferred to 10 ml columns (Poly-Prep Chromatography Column, Bio-Rad) and eluted with LFB1/50 supplemented with 250 mM imidazole.

To outcompete endogenous Ubxn7, recombinant Ubxn7 was used at 0.3 mg/ml in egg extract, while to rescue Ubxn7 depleted extract it was added at 10 μg/ml.

Full length 6xHIS-Faf1 was expressed from pET28a-Faf1 in Rosetta (DE3) pLysS Competent Cells (Novagen, Merck Millipore) bacteria as above. It was purified as Ubxn7 but using the following buffers: lysis buffer (500 mM NaCl, 50 mM Tris-HCl pH 8.0, 2 mM MgCl_2_, 10% glycerol, 0.1% Triton, 1 μg/ml of each: aprotinin, leupeptin and pepstatin, 1 mg/ml lysozyme). After incubation, beads were washed with lysis buffer supplemented with 20 mM imidazole. Beads were eluted with lysis buffer supplemented with 250 mM imidazole.

*Xenopus laevis* GINS was expressed in BL21 (DE3) Competent E coli Cells (C2527H, New England Biolabs) bacteria. The isolated cell pellets were then resuspended into 50 mM Tris pH 7.4, 500 mM NaCl, 0.1 mM PMSF (Sigma) and protease inhibitor tablets (11836170001, Roche). Samples were sonicated for 3X 1 min, 2 sec pulses − 50% duty cycle and supplemented with 1 mg/ml lysozyme. The suspension was then incubated on rocker at 4°C for 1 hour and then clarified by centrifugation at 20,000 g for 30 min at 4°C followed by filtration through 0.22 μm PES filters (Millipore). The supernatant was then passed through 5 ml His-Trap HP column (Cytivia) connected to AKTAprime plus (GE). Gradient elutions were performed with imidazole containing buffer (50 mM Tris pH 7.4, 500 mM NaCl, 500 mM Imidazole and 0.1 mM PMSF). *X. laevis* GINS fractions were eluted with ~79-134 mM imidazole. All fractions were pooled and stored in −80°C.

### Antibodies

α-PCNA (P8825) and α-HIS (H1029) were purchased from Sigma; α-Cul2 (EPR3104) was purchased from Abcam; α-p97 (65278) was purchased from Progen Biotechnik, anti-P-Chk1 (S345) from Cell Signalling.

Affinity purified α-Cdc45, α-Psf2 (12), α-Mcm3 (13), α-LRR1 (S962D) and α-Cul2 (SA206) (4) and α-Mcm7 (6) were previously described. Xenopus Ufd1 antibody was a kind gift from Prof Stemmann’s lab (14).

*Xenopus* full-length p97, Ubxn7, Faf1 and GINS proteins were purified as described above and previously (6,14) and antibodies raised against such prepared antigens in sheep (p97, GINS, Ubxn7) or rabbit (Faf1). The resulting antibody sera was purified in-house against the purified antigen. The specificity of each new antibody is presented in Supplementary Figure 7.

### DNA synthesis assay

Interphase Xenopus *laevis* egg extract was supplemented with 10 ng/μl of demembranated sperm nuclei and incubated at 23°C for indicated time. Synthesis of nascent DNA was then measured by quantification of α^32^P-dATP incorporation into newly synthesised DNA, as described before (15).

### Chromatin isolation time-course

Interphase *Xenopus laevis* egg extract was supplemented with 10-15 ng/μl of demembranated sperm DNA and subjected to indicated treatments. The reaction was incubated at 23°C for indicated length of time when chromatin was isolated in ANIB100 buffer (50 mM HEPES pH 7.6, 100 mM KOAc, 10 mM MgOAc, 2.5 mM Mg-ATP, 0.5 mM spermidine, 0.3 mM spermine, 1 μg/ml of each aprotinin, leupeptin and pepstatin, 25 mM β-glycerophosphate, 0.1 mM Na_3_VO_4_,0.2 μM microcystin-LR and 10 mM 2-chloroacetamide (Merck)) as described previously (15).

During the chromatin isolation procedure, a sample without addition of sperm DNA (no DNA) is processed in an analogous way, usually at the end of the time course, to serve as a chromatin specificity control. The bottom of the PAGE gel on which the chromatin samples were resolved is cut off and stained with Colloidal Coomassie (SimplyBlue, Life Technologies) to stain histones which provide loading controls and indications of sample contamination with egg extract (cytoplasm).

### Nuclei isolation for Chk1 phosphorylation

The nuclei isolation was performed as previously described (5).

### Immunoprecipitation from egg extract

20 μl of egg extract per IP was induced into interphase, and the extract was then supplemented with 4 vol of LFB1/50 buffer (10% sucrose, 50 mM KCl, 40 mM HEPES pH 8, 20 mM K phosphate pH 8, 2 mM MgCl_2_, 1 mM EGTA, 2 mM DTT, 1 μg/ml of each: aprotinin, leupeptin and pepstatin). The diluted extract was cleared of insoluble material by 15 min centrifugation in a microfuge at 4°C, 16k rcf. 100 μl of diluted extract was supplemented with 1 μg of affinity purified Ubxn7, p97 or IgG from sheep serum (15131, Sigma) and incubated on ice for 1h with sporadic mixing. 20 ul of prewashed Protein G Dynabeads (10004D, Life Technologies) were added to each IP sample and incubated for 1 hour at 4°C with rotation. After incubation the beads were washed 3x with LFB1/50 and boiled in NuPAGE LDS loading buffer (Life Technologies).

### Immunoprecipitation from chromatin

100 μl of egg extract per IP was induced into interphase and mixed with 10-15 ng/μl demembranated sperm nuclei and optionally supplemented with the indicated treatments. The reaction was incubated at 23°C for the indicated time. Chromatin was isolated in ANIB100 (50 mM HEPES pH 7.6, 100 mM KOAc, 10 mM MgOAc, 2.5 mM Mg-ATP, 0.5 mM spermidine, 0.3 mM spermine, 1 μg/ml of each aprotinin, leupeptin and pepstatin, 25 mM β-glycerophosphate, 0.1 mM Na_3_VO_4_, 0.2 μM microcystin-LR and 10 mM 2-chloroacetamide), and the chromatin pellets re-suspended in the same volume of original extract of ANIB100 containing 20% sucrose. Protein complexes were released from chromatin by digestion with 2 U/μl of Benzonase nuclease (E1014-25KU, Sigma) and sonicated for 5 min using a Diagenode sonicator with settings: 15s on, 15s off, medium setting. The insoluble fraction was then spun in a microfuge at 4°C, 10 min, 16k rcf.

Prepared beads:

- 30 μl of Dynabeads M-270 epoxy (14302D, Life Technologies) coupled covalently to 20 μg of affinity purified p97 antibody, affinity purified Ubxn7 antibodies or IgG from sheep serum (15131, Sigma);
- 30 μl of Dynabeads Protein G (10004D, Life Technologies) covalently coupled to 6 μg of affinity purified p97 antibody, affinity purified Ubxn7, affinity purified Cul2 or IgG from sheep serum (15131, Sigma) using BS3 crosslinker (S5799, Sigma);

were incubated with 100 μl digested chromatin at 4°C for 1-2 hours with rotation. Following the incubation time, beads were washed for 5 min rotating at 4°C twice with ANIB100, once with ANIB100 containing an additional 0.1% Triton X-100 and finally twice with ANIB100 buffer. Each sample was prepared by boiling in 30 μl of 2x NuPAGE LDS loading buffer (Life Technologies) for 5 min.

### Immunoprecipitation of p97 for mass spectrometry

3.75 ml of *Xenopus laevis* egg extract was activated and supplemented with 10 ng/μl of demembranated sperm DNA, 50 μM p97 inhibitor NMS873 and incubated at 23°C for 60 min. Chromatin was isolated in ANIB/100 buffer. Immunoprecipitation of p97 was performed as described previously (4) and the immunoprecipitated material was analysed by mass spectrometry with Dr Richard Jones from MS Bioworks LLC as previously described (4). The mass spectrometry proteomics data have been deposited to the ProteomeXchange Consortium via the PRIDE repository with the dataset identifier PXD030426 and 10.6019/PXD030426.

### Immunodepletion

Ubxn7 immunodepletions were performed using Dynabeads Protein G (10004D, Life Technologies) coupled to antibodies against Ubxn7 or nonspecific sheep IgGs (15131, Sigma), with two rounds of 1 h incubation at 4°C. The Ubxn7 antibodies were coupled at 600 μg per 1 ml of beads. Effective immunodepletion required 2 rounds of 1 h incubation of egg extract with antibody coupled beads at 50% beads ratio.

Faf1 immunodepletions were performed using Dynabeads Protein A (10002D, Life Technologies) coupled *to Xenopus* Faf1 antibodies raised in rabbit and affinity purified or nonspecific rabbit IgG (15006, Sigma). The Faf1 antibodies were coupled at 600 μg per 1 ml of beads. Effective immunodepletion required 3 rounds of 40 min incubation of egg extract with antibody coupled beads at 50% beads ratio.

### HIS-pulldown from egg extract

30 μl of interphase egg extract per pulldown was supplemented with 10 ng/μl of demembranated sperm nuclei and optionally supplemented with LFB1/50 buffer, or 0.3 mg/ml of recombinant Ubxn7^ΔUBX^, Ubxn7^ΔUIM^ or Ubxn7^wt^ proteins. The replication reaction was stopped with LFB1/50 Buffer supplemented with 0.1% Triton X-100 and chloroacetamide in the middle of the S-phase. The samples were sonicated for 5 min using the Diagenode cold water sonicator with settings: 15s on, 15s off, medium settings. Insoluble material was clarified for 10 min at 4°C at 16k rcf and incubated with 60 μl Dynabeads HIS-Tag isolation (10104D, Invitrogen) for 2 hour with rotation at 4°C. Beads were subsequently washed 2 times with LFB1/50 buffer supplemented with 0.1% Triton X-100 and chloroacetamide. Pulled-down HIS-tagged proteins were eluted by boiling the beads in 30 μl of 2x NuPAGE LDS loading buffer for 5 min.

### Western blot quantification

The quantification of western blots is provided to indicate reproducibility of trends in experiments rather than to provide absolute values of increases or decreases in a signal. The quantified experiments were performed in different preparations of extracts and independently immunodepleted extracts to confirm that observed phenotypes are not specific for one extract prep. As a result, the extracts differ slightly in their kinetics of DNA replication reaction which can affect the levels of detected proteins on chromatin at the same timepoints between different experiments. They do however all reproducibly show the same trend of change across the experiments.

The density of pixels of each band of the western blot and scanned stained histones within the gel were quantified using Image J software. The numeric value in arbitrary units for each band was normalised to loading control (bands of Coomassie stained histones). The analysis of control and treatment samples was always done together and the fold difference between them calculated. Fold change from a number of repeated experiments is plotted on the graphs with mean value and standard error of the mean (SEM) as calculated by GraphPad PRISM.

## Results

### Identification of p97 cofactors acting during unloading of replicative helicase

Using the *Xenopus laevis* egg extract model system, we have previously shown, that the unloading of terminated replicative helicases during S-phase depends on formation of K48-linked ubiquitin chains on the Mcm7 subunit of the CMG helicase by Cul2^Lrr1^ ubiquitin ligase (5). Such modified Mcm7 is in turn recognised and unfolded by p97. We therefore first confirmed that p97 interacts with replicating chromatin in *Xenopus laevis egg* extract with kinetics similar to replication fork components (Figure 1A). We found that p97 is a highly abundant protein in *Xenopus egg* extract (14) and only a small proportion of it interacts with chromatin during DNA replication, p97 starts interacting with chromatin during the exponential stage of replication, when large numbers of replication forks are moving through the chromatin, and is partially retained on chromatin for a while longer than the replicative helicase (Figure 1A). To determine the portfolio of p97 cofactors that interact with p97 during DNA replication termination in egg extract and which may direct p97 to the terminated replisome, we aimed to immunoprecipitate p97 from a chromatin fraction and analyse its interactors. Firstly, we inhibited replisome disassembly by inhibition of p97 activity. Critically, this treatment does not stop p97 from interacting with substrates or chromatin (Supp Fig 1A) and should stabilise p97/substrate complexes on chromatin as the substrates cannot be processed. Subsequently, we isolated chromatin with accumulated terminated replisomes, immunoprecipitated p97 and analysed interacting factors by mass spectrometry. Such analysis revealed numerous components of replication machinery interacting with p97, including those which reside in the replisome built around the CMG helicase, but also other DNA replication and DNA damage repair factors (Supplementary Table 1). In order to focus our analysis on potential p97 cofactors, which direct p97 specifically to the terminated replisomes, results from the p97 interactome were compared with those of an Mcm3 IP, which was performed in conditions blocking replisome disassembly. Briefly, the extract was replicated in the presence of a dominant negative ATPase-dead mutant of p97, described previously, and Mcm3 was immunoprecipitated to isolate terminating replisomes (4,14). This comparison allowed us to determine which of the p97 cofactors identified in the p97 immunoprecipitation are also interacting with the terminated replisomes, as we appreciate that p97 does have other substrates on replicating chromatin (Figure 1B). In doing this, we identified both major cofactors Ufd1 and Np14, which are known to facilitate chromatin functions of p97 segregase and were shown previously by us and others to act in replisome disassembly during S-phase (4,16). Interestingly, only two minor cofactors were identified to interact with both the replisome and p97: Faf1 and Ubxn7 (human UBXD7). Both of these factors have been shown previously to interact preferentially with p97 when in complex with Ufd1/Npl4 (17). To support this finding, we first confirmed by p97 IP and western blotting that p97 can indeed interact with Ubxn7 and Faf1 on S-phase chromatin when replisome disassembly is blocked (Figure 1C).

**Figure 1.**
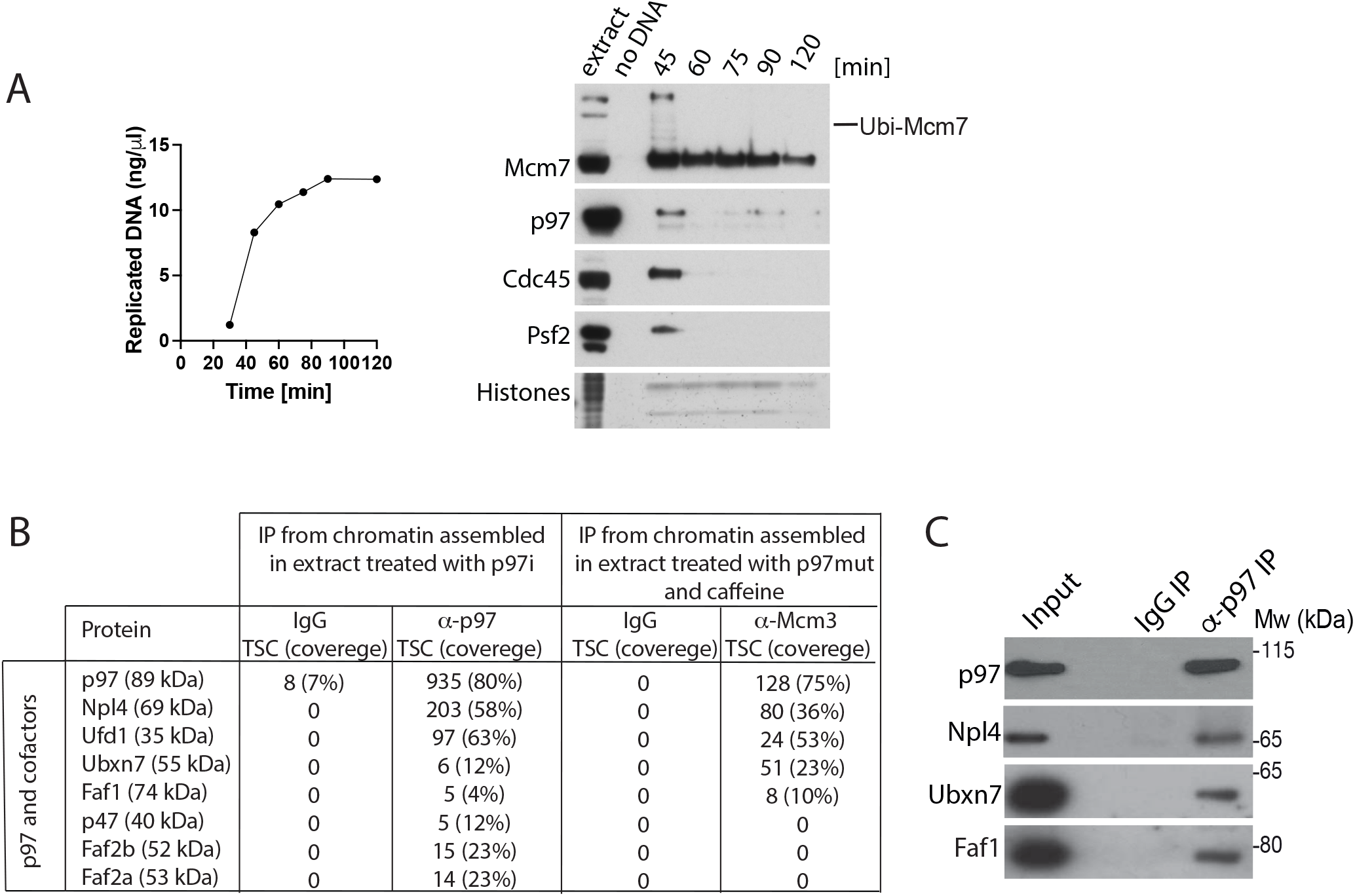
p97 is recruited to chromatin through cofactors. **(A)** Chromatin binding of p97 follows replication fork components. A replication reaction was set up in *X. laevis egg* extract and synthesis of nascent DNA was followed by incorporation of radioactive α-P^32^dATP into newly synthesised DNA (left). At the same time, chromatin was isolated during the replication reaction at indicated time points after sperm DNA addition. Sample without DNA addition was processed in parallel to provide a chromatin specificity control. Histones at the bottom of the PAGE gel were stained with Colloidal Coomassie for loading and sample purity control. Chromatin samples were analysed by western blotting with indicated antibodies (right). **(B)** Ubxn7 and Faf1 are identified as cofactors interacting with both p97 and terminated replisome. Interphase egg extract was supplemented with p97i (NMS873) and chromatin was isolated in late S-phase when high levels of terminated replisomes are accumulated on chromatin. Protein complexes were released from chromatin by Benzonase treatment and proteins interacting with p97 segregase were analysed by mass spectrometry. Identified putative interactors were screened to find known and potential p97 cofactors. Obtained data was compared with the mass spectrometry results of interactors of terminated replisomes published previously (4). Total spectral count is presented with protein coverage in the brackets. Only p97 and its cofactors are presented. **(C)** p97 interacts on chromatin with Ubxn7 and Faf1. A small proportion of input and immunoprecipitated sample from (C) was analysed by western blotting with indicated antibodies.

The finding of Faf1 was somehow unsurprising as it is already known to play a role in maintaining replication fork stability in *C. elegans* and human cell lines (18), and indeed the *C. elegans* homologue of Faf1 (UBXN-3) is essential for replisome unloading in S-phase and in mitotic prophase (4,11). In contrast, this is the first time that Ubxn7 has been implicated in the process of replisome unloading. What is already known about Ubxn7 is that inhibition of the human homologue UBXD7 in human cells leads to hyper-accumulation of DNA damage sensors after UV damage (19), although it is best known as a regulator of degradation of the hypoxia inducible factor Hif1α (20). Interestingly, while targeting Hif1α for degradation, UBXD7 simultaneously interacts with active (neddylated) cullin ligase Cul2^VHL^ through its UIM domain and with p97 through its UBX domain (Supp Fig 2A and B). Given that the same factors appear to be involved in replisome disassembly, we hypothesised that the association of p97 with its cofactor Ubxn7 could provide a mechanism by which p97 targets Mcm7, ubiquitylated with K48-linked ubiquitin chains, synthesised by Cul2^Lrr1^.

### Ubxn7 stimulates efficient replisome disassembly

To investigate the importance of Faf1 and Ubxn7 for replisome disassembly during S-phase, we decided to immunodeplete each protein independently from the egg extract and determine the consequences for replisome unloading. To do this, we have raised antibodies against *X. laevis* Ubxn7 and Faf1 (Supp Fig 7). Both Faf1 and Ubxn7 could be efficiently immunodepleted from egg extracts to less than 5% of total protein (Figure 2A and B). Neither depletion inhibited the extracts’ ability to synthesise nascent DNA in a number of independent immunodepletions (Figure 2A and B), suggesting that neither are essential for DNA synthesis completion in the egg extract. Importantly, neither Ubxn7 nor Faf1 immunodepletion affected the level of each other (Supp Fig 3A). We then followed proteins on chromatin during a replication reaction in IgG-depleted and Ubxn7- or Faf1-depleted extracts. Interestingly, while immunodepletion of Faf1 had a very minor effect on replisome unloading during S-phase (Figure 2D,E), immunodepletion of Ubxn7 reproducibly delayed unloading of replisomes (Cdc45, Psf2) in independently immunodepleted extracts (Figure 2C,E), suggesting that although Ubxn7 is not essential for replisome disassembly, it does regulate the efficiency of this process.

**Figure 2.**
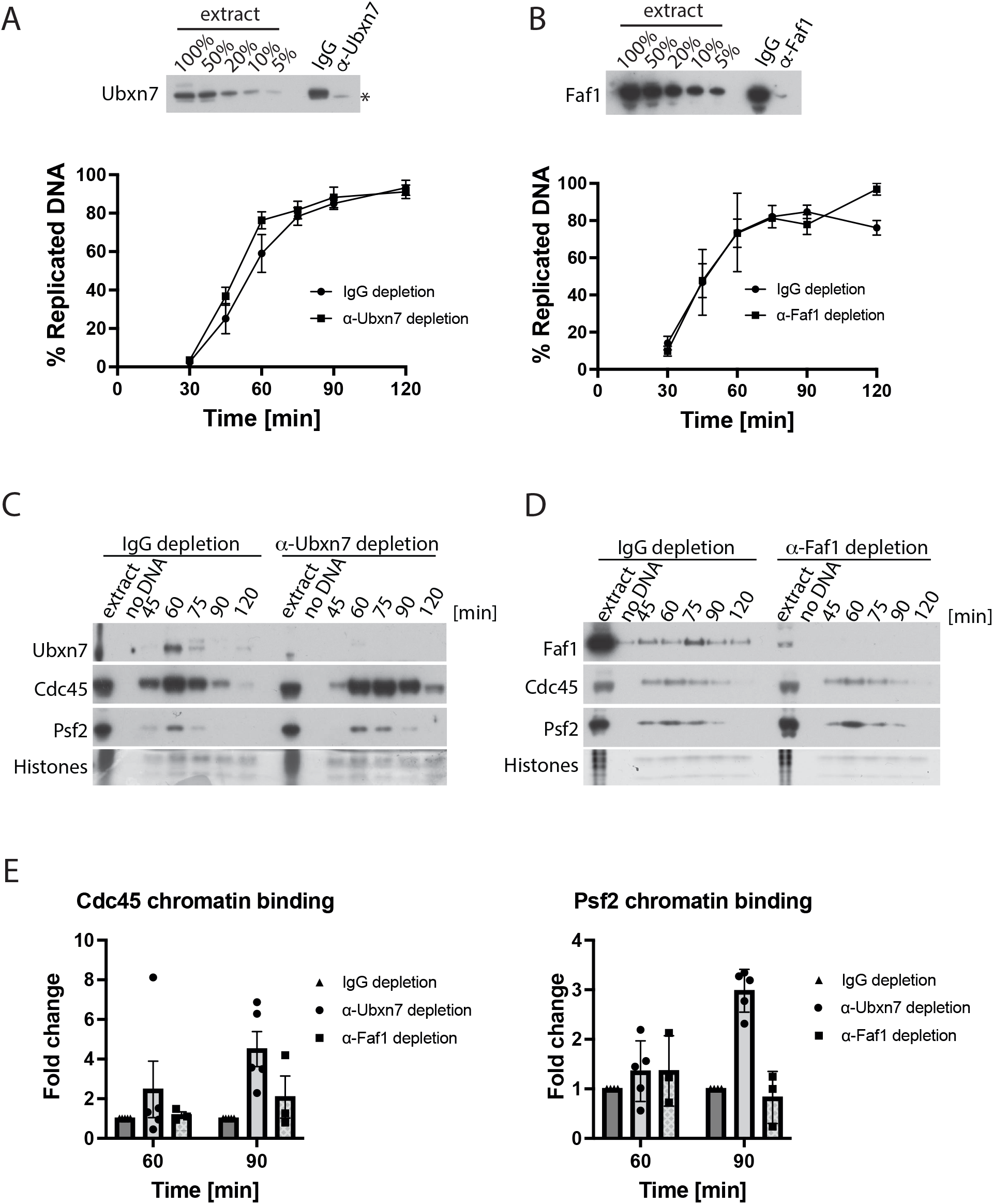
Ubxn7 facilitates terminated replisome unloading. **(A)** Ubxn7 is not required for DNA replication. The remaining level of Ubxn7 upon Ubxn7 immunodepletion was analysed by western blotting. The asterisk indicates a non-specific band (top). The ability of Ubxn7-immunodepleted extract to synthesise nascent DNA was analysed by incorporation of radioactive α-P^32^dATP into newly synthesised DNA (n=6) (bottom). **(B)** Faf1 is not required for DNA replication. The remaining level of Faf1 upon Faf1 immunodepletion was analysed by western blotting (top). The ability of Faf1-immunodepleted extract to synthesise nascent DNA was analysed by incorporation of radioactive α-P^32^dATP into newly synthesised DNA (n=3) (bottom). **(C)** Ubxn7 depletion delays terminated replisomes disassembly. Chromatin was isolated during the replication reaction time course in IgG depleted and Ubxn7 depleted extract. Chromatin samples were analysed as in Figure 1A with indicated antibodies. **(D)** Faf1 depletion does not impact terminated replisomes disassembly. Chromatin was isolated during the replication reaction time course in IgG depleted and Faf1 depleted extract. Chromatin samples were analysed as in Figure 1A with indicated antibodies. **(E)** The fold increase of Cdc45 and Psf2 signal on chromatin in Ubxn7 or Faf1 depleted extracts at indicated time points was quantified in comparison to IgG depletion. For Ubxn7 depleted extract n=5 for Faf1 depleted extract n=3. The mean value is presented with all individual points and with standard error of the mean (SEM) as error bars.

### Importance of Ubxn7 for replisome disassembly

Next, we wanted to understand how the depletion of Ubxn7 delays the unloading of the replisomes in the egg extract. Induction of replication stress, which affects progression of replication forks, can lead to transient accumulation of replisomes on chromatin. However, Ubxn7 depletion did not lead to the accumulation of DNA damage-associated replication stress, as determined by γH2AX immunoblotting (Figure 3A), nor the phosphorylation of Chk1 (Supp Fig 3B). Taken together, this demonstrates that accumulation of replisomes on chromatin in Ubxn7-depleted extract is not due to the effects of replication stress.

**Figure 3.**
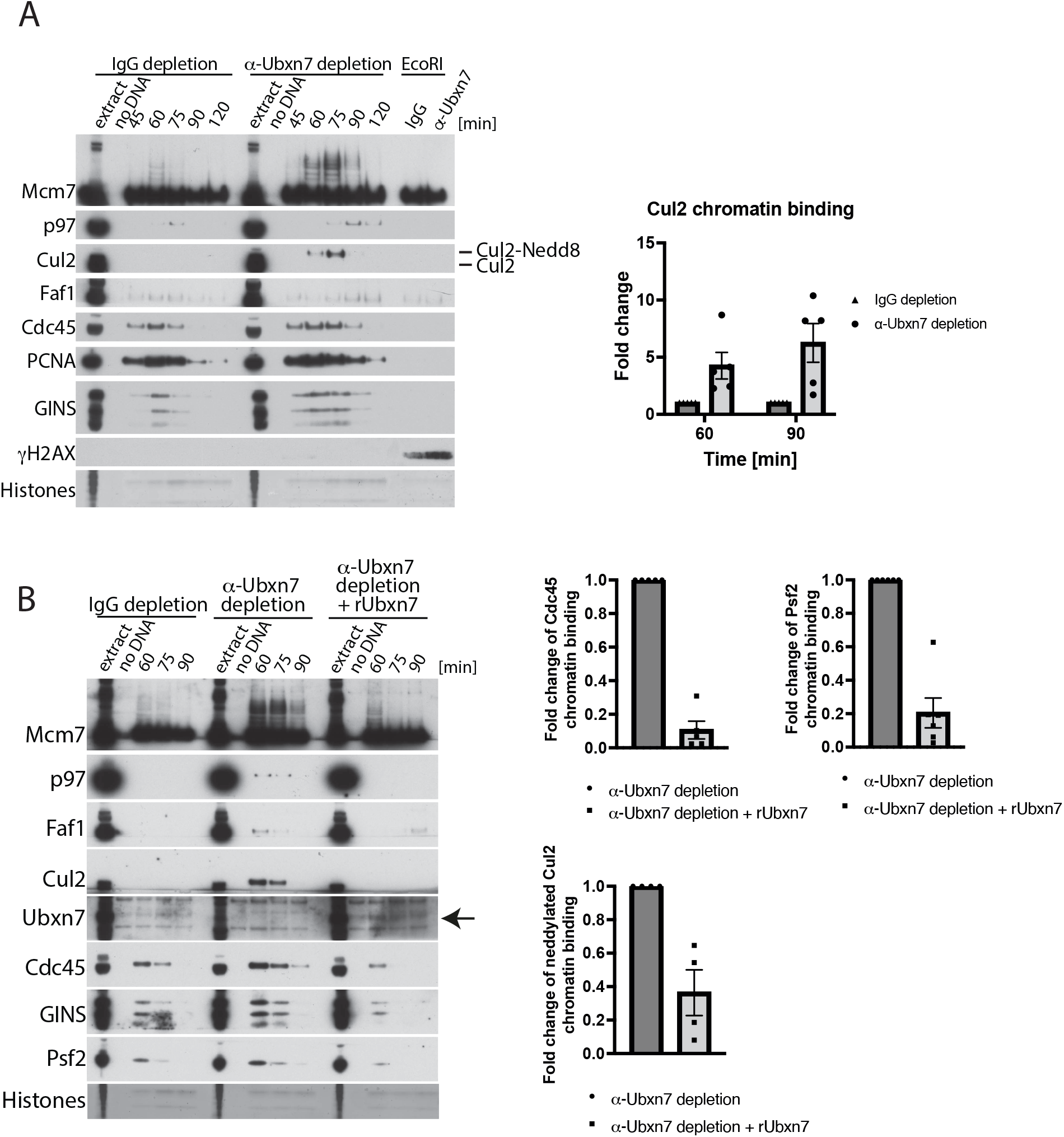
Ubxn7 depletion leads to accumulation of ubiquitylated Mcm7 and active Cul2^Lrr1^ on chromatin. **(A)** Chromatin was isolated during the replication reaction time course in IgG-depleted and Ubxn7-depleted extract. Chromatin samples were analysed as in Figure 1A with indicated antibodies. The fold increase of Cul2 signal on chromatin in Ubxn7 depleted extract at indicated time points was quantified in comparison to IgG depletion (n=5). The mean value is presented with all individual points and with standard error of the mean (SEM) as error bars. **(B)** Chromatin was isolated during the replication reaction time course in IgG-depleted, Ubxn7-depleted and Ubxn7-depleted extract supplemented with recombinant Ubxn7 at 10 μg/ml. Samples analysed as in (A). The rescue of the phenotypes by rUbxn7 was quantified in 4 experiments and the fold downregulation of CMG components and Cul2 retainment presented as a mean value with SEM.

We next analysed how Ubxn7 depletion affects the levels of Mcm7 ubiquitylation and chromatin-bound Cul2 and p97 (Figure 3A). Interestingly, while the levels of p97 and Faf1 on chromatin were slightly increased by Ubxn7 depletion (Figure 3A), Cul2 markedly accumulated on chromatin in its active neddylated form (Figure 3A, neddylated Cul2 runs at a higher molecular size on the gel – compare size of unneddylated Cul2 in egg extract and neddylated on chromatin). Similarly, ubiquitylated Mcm7 also accumulated on chromatin, modified with longer ubiquitin chains.

We then decided to ensure that these phenotypes are actually caused by immunodepletion of Ubxn7 rather than an unidentified protein recognised by Ubxn7 antibody, through supplementing immunodepleted extract with recombinant Ubxn7. As shown in Figure 3B, addition of recombinant Ubxn7 could rescue the delayed unloading of CMG components (Cdc45 and Psf2), the accumulation of Cul2 on chromatin and the accumulation of long-chain modified Mcm7 (Figure 3B). These results, suggest that Ubxn7 may be fine-tuning the process of replisome unloading, acting as a bridge between Cul2^Lrr1^, p97 and their shared substrate Mcm7. This bridging allows for fast and efficient processing of Mcm7 ubiquitylated with relatively short ubiquitin chains. Without Ubxn7, Cul2^Lrr1^ recognises terminated CMG and ubiquitylates Mcm7, but this ubiquitylated Mcm7 is not recognised and processed quickly by p97 due to lack of Ubxn7. Cul2^Lrr1^ stays therefore associated with Mcm7 for a longer time (we can observe this as accumulation of active Cul2 on chromatin), resulting in synthesis of longer ubiquitin chains on Mcm7. Although this most likely enables eventual recruitment of p97, the process is less efficient.

### Regulation of Ubxn7 during DNA replication in egg extract

To determine how Ubxn7 is regulated during replication termination and replisome disassembly in the egg extract, we analysed its pattern of chromatin binding during a replication reaction in egg extract. Throughout normal, unchallenged replication, Ubxn7 transiently interacts with chromatin with timing concomitant to that of replication fork presence and Cul2^Lrr1^ (Figure 4A and B (DMSO control)). Inhibition of replisome unloading with p97 ATPase inhibitor led to an accumulation of not only CMG and Cul2^Lrr1^ on chromatin as expected (4), but also of Ubxn7 and p97 (Figure 4A and Supp Fig 4A). In contrast, while inhibition of cullin activity with neddylation inhibitor MLN4924 led to accumulation on chromatin of CMG and Cul2^Lrr1^ as we reported previously (4), levels of chromatin-bound p97 and Ubxn7 were reduced (Figure 4B and Supp Fig 4B). This result suggests that the key determinants of p97 chromatin binding during DNA replication in the egg extract are either cullin-driven ubiquitylation of substrates and/or neddylation of cullins.

**Figure 4.**
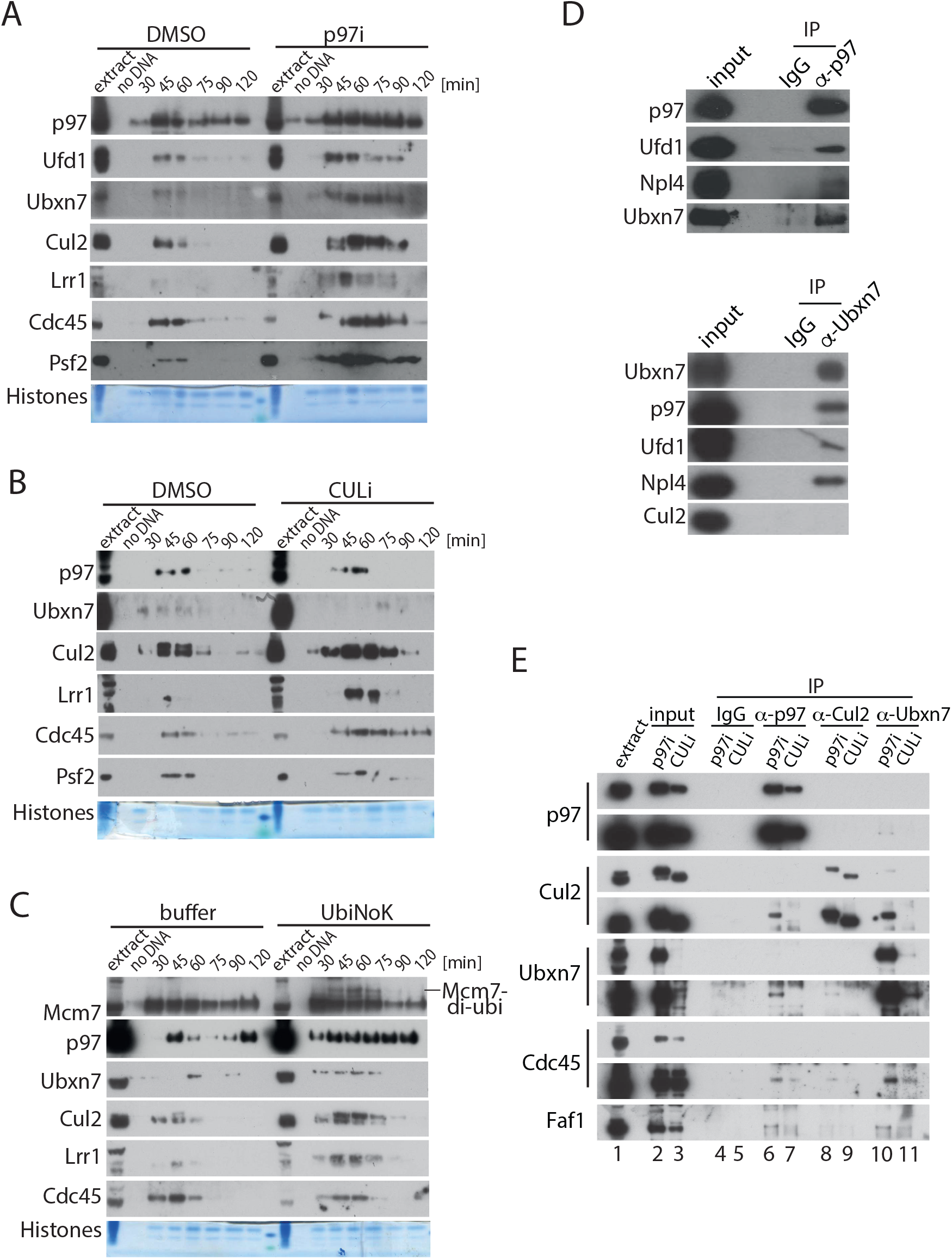
Regulation of Ubxn7 chromatin binding. **(A)** p97 and Ubxn7 accumulate on chromatin upon p97 activity inhibition. Interphase egg extract was supplemented with DMSO or p97i and chromatin was isolated during the replication reaction. Chromatin samples were analysed as in Figure 1A and quantified in Supp Fig 4A. **(B)** p97 and Ubxn7 decrease on chromatin upon Cullin activity inhibition. Interphase egg extract was supplemented with DMSO or CULi and chromatin was isolated during the replication reaction. Chromatin samples were analysed as in (A) and quantified in Supp Fig 4B. **(C)** p97 and Ubxn7 accumulate on chromatin upon inhibition of polyubiquitylation. Interphase egg extract was supplemented with LFB1/50 buffer or 6HIS-UbiNoK and chromatin was isolated during the replication reaction. Chromatin samples were analysed as in (A) and quantified in Supp Fig 4C. **(D)** Ubxn7 interacts with p97 but not with Cul2 in egg extract. Ubxn7 or p97 were immunoprecipitated from egg extract. Interacting partners were analysed by western blotting with indicated antibodies. **(E)** Interaction between Ubxn7, p97 and Cul2 is disrupted when neddylation of Cul2 is inhibited. Interphase egg extract was supplemented with CULi (MLN4924) or p97i (NMS873). Chromatin was isolated in late S-phase when a high level of post-termination replisomes accumulated on chromatin, protein complexes were released from chromatin by Benzonase treatment and p97, Cul2 or Ubxn7 immunoprecipitated from the chromatin proteome. Immunoprecipitated samples were analysed by western blotting with indicated antibodies. Short and long exposures for each of the immunoprecipitated proteins is presented.

Finally, we decided to inhibit the polyubiquitylation of all the potential substrates of p97 during DNA replication by supplementing the extract with a chain terminating mutant of ubiquitin: 6xHIS-UbiNoK. Interestingly, despite inhibition of polyubiquitylation by6xHIS-UbiNoK, as observed through accumulation of di-monoubiquitylated forms of Mcm7 on chromatin, we can reproducibly observe a higher level of p97 and Ubxn7 binding to chromatin (Figure 4C and Supp Fig 4C). Importantly, neither inhibition of cullins nor global polyubiquitylation affects the extracts’ ability to replicate DNA (Supp Fig 4B and C). Our findings therefore suggest that Ubxn7 behaves like a p97 cofactor and follows the same patterns of chromatin interaction. Moreover, p97 segregase is not directed to chromatin and terminated replisomes just simply through interaction with polyubiquitylated substrates, as inhibition of polyubiquitylation does not prevent p97 and Ubxn7 from binding to chromatin (Figure 4C and Supp Fig 4C). It is likely therefore that treatment of replicating extract with the cullin neddylation inhibitor (MLN4924) (Figure 4B and Supp Fig 4B) downregulates chromatin binding of p97 and Ubxn7 because neddylated cullins provide a binding platform for Ubxn7.

To support this hypothesis further, we confirmed that Ubxn7 can interact with p97 not only on chromatin (Figure 1C) but also in the egg extract (cytoplasm) (Figure 4D). However, we could not detect an interaction between Ubxn7 and Cul2 in the egg extract (Figure 4D), as Cul2 is present in the cytoplasmic extract in its inactive, unneddylated form. We went on to perform reciprocal immunoprecipitations of p97, Cul2 and Ubxn7 from chromatin where post-termination replisomes were accumulated due to inhibition of Cullin neddylation (CULi) or p97 activity (p97i) (Figure 4E). The level of inhibition achieved in the samples can be judged by the status of Cul2 in the input: while Cul2 is present in the extract mainly in its inactive, unneddylated form (Figure 4E, lane 1), it accumulates on chromatin during termination upon inhibition of p97 activity mostly in its active, neddylated form (Figure 4E, lane 2). CULi treatment, however, leads to Cul2 accumulation on chromatin in its inactive unneddylated form (Figure 4E, lane 3). The dramatic absence of Ubxn7 on chromatin upon neddylation inhibition, despite the presence of p97 and Faf1 in this input (Figure 4E, lane 3), clearly suggests that Ubxn7 binding to chromatin strongly depends on Cullin neddylation. When present in the chromatin input, Ubxn7 could co-IP neddylated Cul2, Faf1 and a little of p97 (Figure 4E, lane 10). p97 could interact with Ubxn7, Faf1 and neddylated Cul2 but this can only be detected when replisomes have accumulated in their ubiquitylated form, due to inhibition of p97 activity (Figure 4E, compare lane 6-7). Similarly, despite immunoprecipitation of equal quantities of neddylated and unneddylated Cul2 from each sample, Cul2 could only interact with Ubxn7 when in its neddylated form on p97i treated chromatin (Figure 4E, lane 8 and 9). Reassuringly, all three: p97, Cul2 and Ubxn7 could co-IP a little of the component of the terminated replisome Cdc45. Altogether, these experiments suggest that Ubxn7, although being a p97 cofactor, is recruited to chromatin during the termination reaction through its interaction with neddylated Cul2. Moreover, Faf1 is likely to form a common complex with p97 and Ubxn7 as we can see it interacting with Ubxn7 when present on chromatin (Figure 4E, lane 10).

### Ubxn7 bridges Cul2^Lrr1^ and p97 through its UIM and UBX domains

Our results suggest that, analogously to Hlf1α regulation, Ubxn7 acts as a bridge between Cul2^Lrr1^, its substrate Mcm7 and the p97 segregase complex. To explore this idea in more detail we decided to make use of separation-of-function mutants of Ubxn7 that cannot interact with p97 (UBX domain mutated, rUbxn7^ΔUBX^) or Cul2 (UIM domain mutated, rUbxn7^ΔUIM^) (Supp Fig 2A and C). In human cells the P459G mutation abolishes interaction with p97, while S297A abolishes interaction with neddylated Cul2, while not affecting p97 interaction (21). Moreover, L290, A293 and S297 were found to be the most conserved aminoacids in UIMs of several human proteins (22). We have therefore mutated corresponding P458G in the *Xenopus laevis* Ubxn7 sequence to create rUbxn7^ΔUBX^ and the corresponding L286E/A289Q/S293A residues to create rUbxn7^ΔUIM^ (Supp Fig 5A). While we were able to confirm that rUbxn7^ΔUBX^ cannot interact with p97 in the egg extract (Supp Fig 5B), Ubxn7 and Cul2 do not interact in the egg extract (cytoplasm) and so it is not easy to verify whether the rUbxn7^ΔUIM^ mutation disrupts this. We did observe, however, that adding a high concentration of recombinant Ubxn7^ΔUIM^ mutant to normal egg extract with endogenous Ubxn7 present (mimicking overexpression experiments), caused a substantial increase in active, neddylated Cul2 on chromatin. This was not observed upon addition of wt rUbxn7 or rUbxn7^ΔUBX^ mutants (Supp Fig 5C). This suggests that outcompeting endogenous Ubxn7 with a mutant that cannot interact with neddylated Cul2 reproduces the phenotype of Ubxn7 immunodepletion i.e. increased and prolonged association of Cul2 with chromatin (Figure 3A). Importantly, addition of neither protein affected the extract’s ability to synthesise nascent DNA (Supp Fig 5D). We are confident therefore that Ubxn7^ΔUIM^ is defective in binding to neddylated Cul2.

To further test functionality of these mutants, Ubxn7 immunodepleted extract was supplemented with either wt or mutant Ubxn7. Neither addition to immunodepleted extract blocked the extract’s ability to perform nascent DNA synthesis (Supp Fig 6A). While addition of wt Ubxn7 could support timely unloading of CMG from chromatin and prevent excessive accumulation of Cul2 on chromatin and long ubiquitin chain formation on Mcm7, neither of the two mutants could fully rescue the Ubxn7 immunodepletion phenotypes (Figure 5A and Supp Fig 6B). This indicates that both domains are important for Ubxn7 function during replisome disassembly. Altogether these data suggest that through its UBX and UIM domains, Ubxn7 can indeed bridge Cul2^Lrr1^ and the p97 complex, and that binding to neddylated Cul2^Lrr1^ through its UIM domain is especially important for restricting Cul2 activity and/or stimulating its dissociation from the terminated replisome during replication termination.

**Figure 5.**
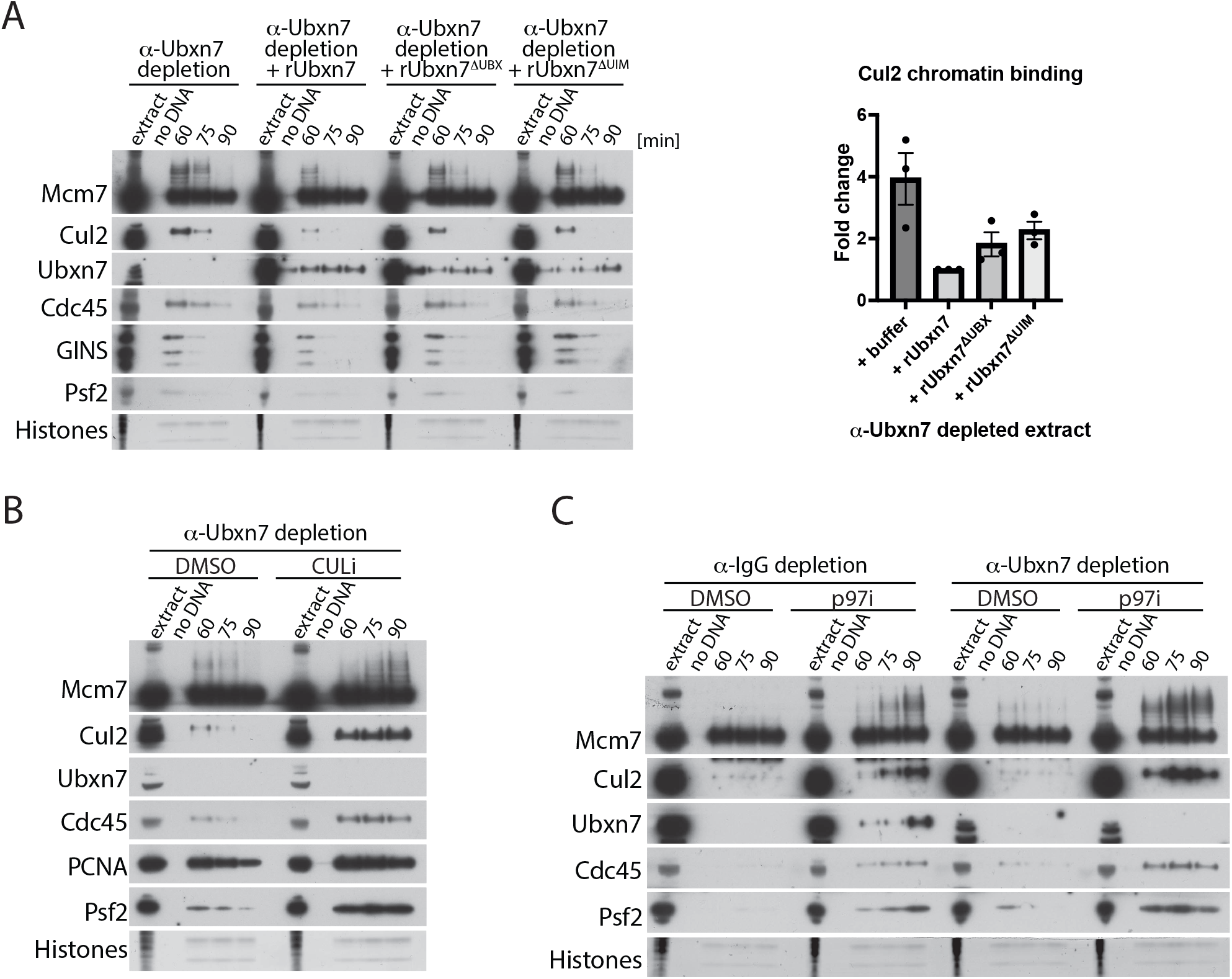
Ubxn7 bridges Cul2^Lrr1^ and p97 complexes leading to efficient unloading of ubiquitylated Mcm7. **(A)** Both UIM and UBX domains of Ubxn7 are important for its functions. Ubxn7 depleted extract was supplemented with recombinant Ubxn7 or point mutants that disrupt UBX or UIM domain functions (Ubxn7^ΔUB×^ and Ubxn7^ΔUIM^, respectively). Chromatin was isolated at indicated time points and analysed as in Figure 1A. The level of Cul2 chromatin binding at 60 min time point was quantified over 3 experiments and the fold rescue of the Cul2 accumulation in comparison to Ubxn7 depleted extract is presented. Individual values, mean and SEM are shown. CMG unloading is quantified in Supp Fig 6B. **(B)** Cullin activity is needed for replisome unloading in absence of Ubxn7. Ubxn7 depleted extract was supplemented with DMSO or CULi and chromatin samples isolated during the replication reaction and analysed as in (A). **(C)** The chains built on Mcm7 in Ubxn7 depleted extract are shorter than those built upon p97 activity inhibition. IgG- or Ubxn7-depleted extracts were optionally supplemented with p97i. Chromatin samples were analysed at indicated time points as in (A).

### Unrestricted Cul2 activity allows for Mcm7 unloading in Ubxn7 depleted extract

Our results above show that upon Ubxn7 depletion, we observe accumulation on chromatin of active, neddylated Cul2^Lrr1^ and ubiquitylated forms of Mcm7. It is likely therefore that continuous growth of the length of chains on Mcm7 finally leads to p97 recognition and unloading. To test that it is Cul2^Lrr1^ and not a different ubiquitin ligase (e.g. TRAIP) synthesising these long chains, we blocked Cul2 activity using CULi in IgG- and Ubxn7-depleted extracts (Figure 5B). Indeed, replisome disassembly was blocked in Ubxn7-depleted extract treated with CULi, and the ubiquitylation of Mcm7 observed in Ubxn7-depleted extract at 60 min was strongly inhibited by CULi. Instead, we observed a much more gradual accumulation of ubiquitylated Mcm7, as we always do upon CULi treatment (Figure 5B, compare also Supp Fig 6C). This shows that Cul2 activity is still required for replisome disassembly in the absence of Ubxn7.

Finally, we wanted to assess whether the length of chains built on Mcm7 upon Ubxn7 depletion is unusually long, suggesting uncontrolled Cul2^Lrr1^ activity or whether it is comparable to the level of ubiquitylation we observe upon blocking p97 segregase activity and replisome unloading. To this end we inhibited p97 in Ubxn7 depleted extract (Figure 5C) and could see that the level of ubiquitylation upon complete inhibition of unloading with p97i is even higher, suggesting that it is just the delay in replisome disassembly that gives Cul2^Lrr1^ more time to ligate longer ubiquitin chains.

Altogether, our data support a model whereby Ubxn7 binds to active, neddylated Cul2^Lrr1^ on chromatin to facilitate fast recruitment of the p97 complex to ubiquitylated replisomes resulting in efficient replisome unloading (Figure 6A). In the absence of Ubxn7, Cul2^Lrr1^ can still bind to terminated replisome but recruitment of p97 is delayed. In the meantime, active, neddylated Cul2^Lrr1^ keeps ubiquitylating Mcm7, forming longer ubiquitin chains, which finally allow for p97 recognition and replisome disassembly (Figure 6B).

**Figure 6.**
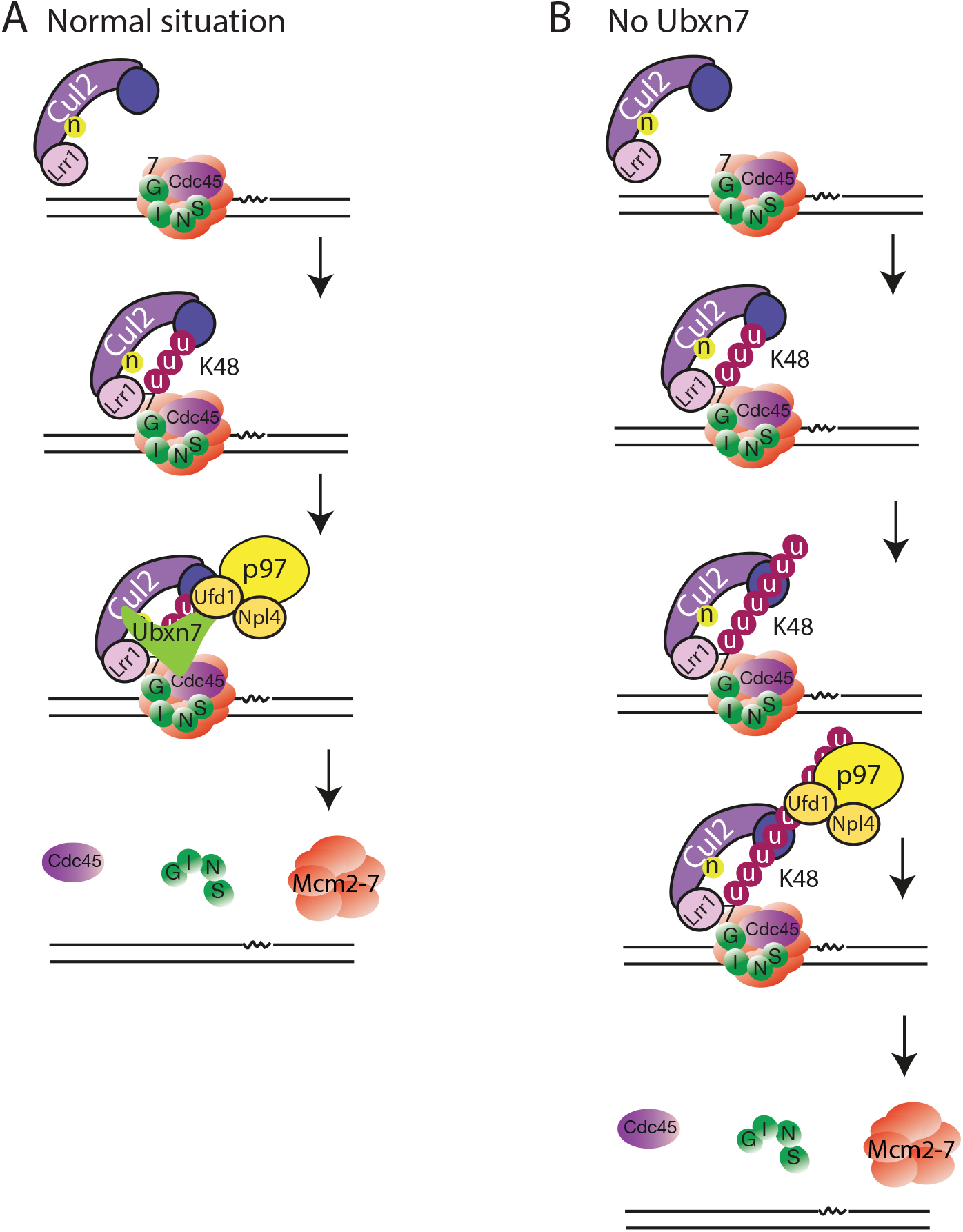
Proposed model of Ubxn7 function. **(A)** Ubxn7 is bridging Cul2^Lrr1^, ubiquitylated Mcm7 and the p97 complex, leading to efficient CMG helicase unloading. **(B)** Delays in replisome unloading upon lack of Ubxn7. p97 can still recognise ubiquitylated Mcm7 but the process is slower and takes longer time.

## DISCUSSION

### Ubxn7 streamlines replisome disassembly during replication termination

Our results suggest that by concomitant interactions with neddylated Cul2^Lrr1^, ubiquitylated Mcm7 and p97 complex, Ubxn7 facilitates efficient and fast unloading of terminated CMG helicases from chromatin (Figure 6). The general mode of Ubxn7 operation in replisome disassembly during termination closely resembles the way UBXD7 regulates degradation of Hif1α in collaboration with CUL2^VHL^ and p97/UFD1/NPL4 (20,21). It is interesting to speculate that Ubxn7 may not only bridge the three factors to facilitate recognition of terminated ubiquitylated replisomes, but may also stimulate turnover of Cul2^Lrr1^ and promote its dissociation from terminated replisomes. This in turn could facilitate the unfolding of ubiquitylated Mcm7 by the p97 complex.

The ability of Ubxn7 to facilitate unloading of Mcm7 ubiquitylated with short ubiquitin chains synthesised by Cul2^Lrr1^ explains also why we observe such a strong accumulation of Mcm7 modified with short ubiquitin chains after treatment of replication reactions with CULi (Figure 5B and Supp Fig 6C). CULi inhibits neddylation of Cullins, so it not only slows down the activity of Cul2^Lrr1^, but also inhibits the interaction of Ubxn7 with Cul2^Lrr1^. As a result, the slowly building up chains on Mcm7 have to reach a higher threshold of length to be efficiently extracted by the p97 complex.

The function of Ubxn7/UBXD7 in streamlining the process of replisome disassembly may not just be through accelerated recognition of ubiquitylated substrate by the p97 segregase complex, but also by increasing the rate of substrate unfolding. A recent study has shown that once the substrate is recognised, p97 starts substrate processing by unfolding one of the distal ubiquitins in the ubiquitin chain attached to the substrate. It then pulls both the unfolded ubiquitin chain and the unfolded substrate through the central channel of the hexamer (23), which leads to extraction of the substrate from complex structures. In the absence of Ubxn7, when Mcm7 is modified with long ubiquitin chains, the process of unfolding of such modified Mcm7 is likely to be slower as it is more likely for p97/Ufd1/Npl4 to bind the ubiquitin chain further away from the Mcm7 substrate, which necessitates unfolding of a longer ubiquitin chain before unfolding Mcm7 itself.

### Ubxn7 and Faf1 during replication

Immunodepletion of Ubxn7, but not Faf1, from egg extract leads to a delay in replisome disassembly (Figures 2 and 3). The *C. elegans* homologue of Faf1, UBXN-3, has been shown to be important for CMG helicase unloading by p97 in S-phase and in mitosis (4,11), but also to regulate other replication factors such as CDT-1 and, recently, to regulate SUMOylated factors at DNA replication forks (18,24,25). It is clear, therefore, that UBXN-3 plays a key role in extraction of proteins from chromatin during DNA replication in *C. elegans* embryos. There is, however, no homologue of Ubxn7 in *C. elegans*, and UBXN-3/Faf1 does not contain a UIM domain that could direct it to neddylated cullins, so it is unlikely to substitute for the role that Ubxn7 plays in vertebrates.

In our experiments, in the absence of Ubxn7, the unloading of the replisomes is delayed but they are still eventually unloaded (Figure 2 and 3). At the same time, we observe accumulation of active Cul2^Lrr1^ and higher levels of ubiquitylated Mcm7 on chromatin (Figure 3). After a delay, p97/Ufd1/Npl4 can, therefore, recognize ubiquitylated Mcm7 and extract it, possibly with help from other cofactors such as Faf1. In other organisms that have no Ubxn7, this process is also likely to be facilitated by other cofactors such as Faf1/UBXN-3. Interestingly, in an *in vitro* reconstitution system of DNA replication with purified budding yeast proteins, where no p97 cofactors were present, Mcm7 needed to be ubiquitylated with a minimum ubiquitin chain length of 5 in order to be recognized and extracted by the CDC48 complex. This process, however, worked much better with longer chains (26). The evolution of Ubxn7 to specifically link neddylated cullins with their substrates and the processing factor p97 is an additional level of regulation that ensures efficiency in p97 substrate targeting.

The role of UBXN-3 in regulating SUMOylated factors at replication forks seems to have been conserved throughout evolution as it is also the case in human immortalized cells (25). It will be very interesting to decipher the contribution of FAF1 and UBXD7 to replisome disassembly in human cells in the future.

### Importance of efficient replisome disassembly

Replisome unloading must be carefully regulated to maintain genome stability. Premature replisome unloading would likely lead to a collapse of replication forks and creation of DNA damage (27). However, defects in timely replisome disassembly are also detrimental for cells. For example, *dia2Δ* budding yeast cells, which cannot ubiquitylate Mcm7 during termination, are defective in cell cycle progression, present high levels of genomic instability, are unable to grow at low temperatures and are sensitive to drugs that compromise replication fork progression (28–31). Genetic loss of *lrr-1* in worms results in mitotically arrested *C.elegans* embryos and germ lines (32,33). Moreover, while partial disruption of the S-phase or mitotic pathways of replisome disassembly alone had no effect on worm embryo viability, disrupting both pathways led to embryonic lethality (4). Finally, CRISPR/Cas9-mediated deletion of LRR1 or TRAIP in a number of human cell lines is lethal (34). Interestingly, a recent study suggested that efficient LRR1-mediated replisome disassembly is essential for completion of DNA replication, possibly through recycling of replisome components from early activated replication forks to late firing ones (35). With all these in mind, it is clear that timely and efficient replisome disassembly is important for the maintenance of genome stability and that Ubxn7 is pivotal for this.

## Supporting information

combined supplemental figures and table

## Acknowledgements

We would like to thank Dr Rebecca Jones, Dr Neville Gilhooly, Dr Marco Saponaro, Dr Paloma Garcia and Dr Clare Davies for critical discussions of the manuscript. We would also like to thank our families for the continuous support.

