## Supplementary material for "The p97 cofactor Ubxn7 facilitates replisome disassembly during S-phase": combined supplemental figures and table

Tarcan et all Supp Figure 1

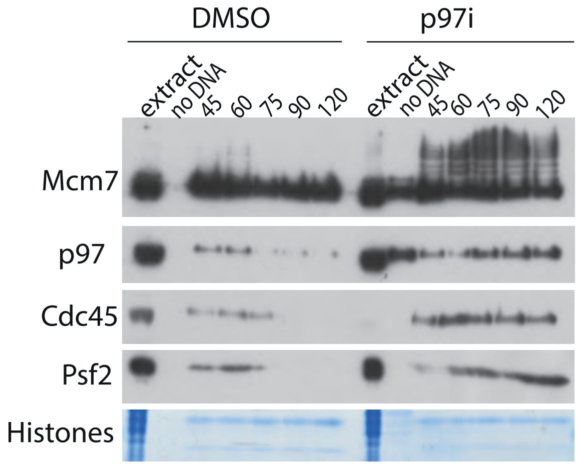


**Supplementary Figure 1**
p97 accumulates on chromatin upon inhibition of its ATPase activity. Interphase egg extract was supplemented with DMSO or p97i and chromatin samples isolated at indicated timepoints during replication reaction. Chromatin samples were analysed by western blotting with indicated antibodies as in Figure 1A. Presence of a ladder of bands in -DNA control in p97i indicates contamination with cytoplasm in this particular sample. See also Figure 4A for alternative western blot. CMG components (Cdc45 and Psf2) unloading is inhibited with p97i and ubiquitylated forms of Mcm7 accumulate on chromatin when p97 is not active.

Tarcan et all Supp Figure 2

A

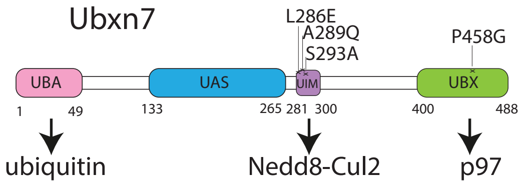


B

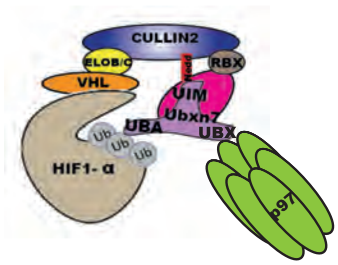


C

```
X.l.   MEKDGG-----VSQQLLREFTAITGATDSVAKHMLEACNHNLEMAVTMFLDGGGIPDEPS     55
H.s.   MAAHGGSAASSALKGLIQQFTTITGASESVGKHMLEACNNNLEMAVTMFLDGGGIAEEPS     60
       *  .**     . : *:::**:****::**.********:*************** :***

X.l.   TSSAGSSTARPAPDLSDDVRAPIPQKQEILVEPE-IFGAPKRRRPARSIFDGFRDFQTET     114
H.s.   TSSASVSTVRP--HTEEEVRAPIPQKQEILVEPEPLFGAPKRRRPARSIFDGFRDFQTET     118
       ****. **.**  . .::**************** :************************

X.l.   IRQEQELRNGGAVDKKLTTLADLFRPPIDLMHKGSFETAKQFGQLHNKWLMINIQNVQDF     174
H.s.   IRQEQELRNGGAIDKKLTTLADLFRPPIDLMHKGSFETAKECGQMQNKWLMINIQNVQDF     178
       ************:***************************: **::**************

X.l.   ACQCLNRDIWSNDAIKTLIREHFIFWQVYHDSEEGQRYIQFYKLPEFPYVSILDPRTGQK     234
H.s.   ACQCLNRDVWSNEAVKNIIREHFIFWQVYHDSEEGQRYIQFYKLGDFPYVSILDPRTGQK     238
       ********:***:*:*.:************************** :**************

X.l.   LVEWHKLDPNSFVEQVTGFLGEHGQLDGLSSSPPKKRLRSESLIDASEDSQLEAAIRASL     294
H.s.   LVEWHQLDVSSFLDQVTGFLGEHGQLDGLSSSPPKKCARSESLIDASEDSQLEAAIRASL     298
       *****:**  .**::**********************  **********************

X.l.   QETHFDSAVNKETPRREEESESELYSGSEEFISVCGSDHEEDEERPVIESSAEPDSCGKS     354
H.s.   QETHFDSTQTKQDSRSDEESESELFSGSEEFISVCGSDEEEEVENLA-----------KS     347
       *******: .*:  * :*******:*************.**: *. .           **

X.l.   KKSTSKDPNHRKEEVGRTH--PVVVEE---TSNHQSPE---DTASSEEHHRESGCSVAVS     406
H.s.   RKSPHKDLGHRKEENRRPLTEPPVRTDPGTATNHQGLPAVDSEILEMPPEKADGVVEGID     407
       :**  ** .*****  *    * *  :   ::***.     .   .   .: .*   .:.

X.l.   DNGPKAKLMLRYPDGKKEQIALPEQAKLLALVRHVQLKGYPNERFELLTNFPRRKLSHLD     466
H.s.   VNGPKAQLMLRYPDGKREQITLPEQAKLLALVKHVQSKGYPNERFELLTNFPRRKLSHLD     467
        *****:*********:***:***********:*** ***********************

X.l.   YDVTLQEAGLCPQETVFVQERN    488
H.s.   YDITLQEAGLCPQETVFVQERN    489
       **:*******************
```

**Supplementary Figure 2**

**(A)** Model of *Xenopus* Ubxn7 structure. The key domains are highlighted together with the mutations incorporated to disrupt UIM and UBX interactions. **(B)** Model of Ubxn7 function during Hif1$\alpha$ processing. **(C)** Comparison of *Xenopus laevis* and human Ubxn7 protein sequence. The domains are highlighted in the same colour as in (A). Amino acids mutated in Ubxn7$^{\Delta UBX}$ and Ubxn7$^{\Delta UIM}$ mutants are highlighted in red.

Tarcan et al Supp Figure 3

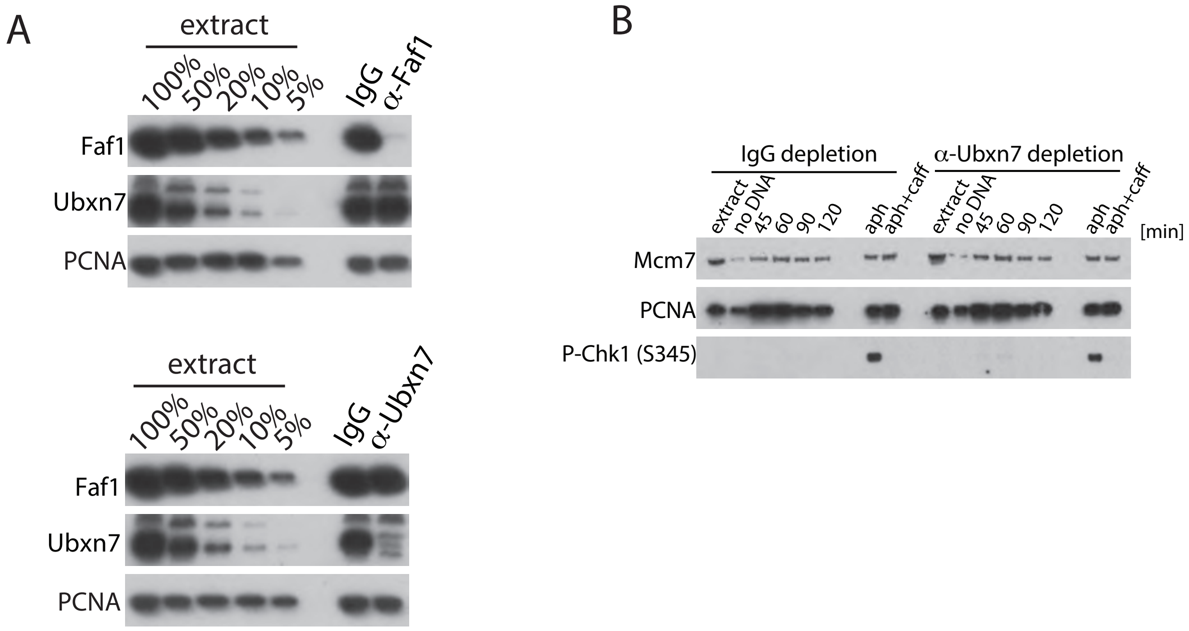


**Supplementary Figure 3**

**(A)** Immunodepletion of Ubxn7 or Faf1 does not co-deplete each other. Faf1 and Ubxn7 were immunodepleted independently as described in materials and methods. The level of remaining proteins in the egg extract was analysed through western blotting of a series of dilutions of egg extract and samples of IgG-, Ubxn7- and Faf1-depleted extracts. PCNA serves as an example of protein that is not affected by either immunodepletion. **(B)** Immunodepletion of Ubxn7 does not lead to checkpoint activation and Chk1 phosphorylation. Nuclei were isolated at indicated timepoints during replication reaction in IgG- or Ubxn7-depleted extracts. Nuclei samples were analysed by western blotting with indicated antibodies. As a positive control a sample of each extract was treated with polymerase inhibitor aphidicolin (polymerase inhibitor which induces checkpoint activation) or aphidicolin and caffeine (checkpoint inhibitor).

Tarcan et al Supp Figure 4

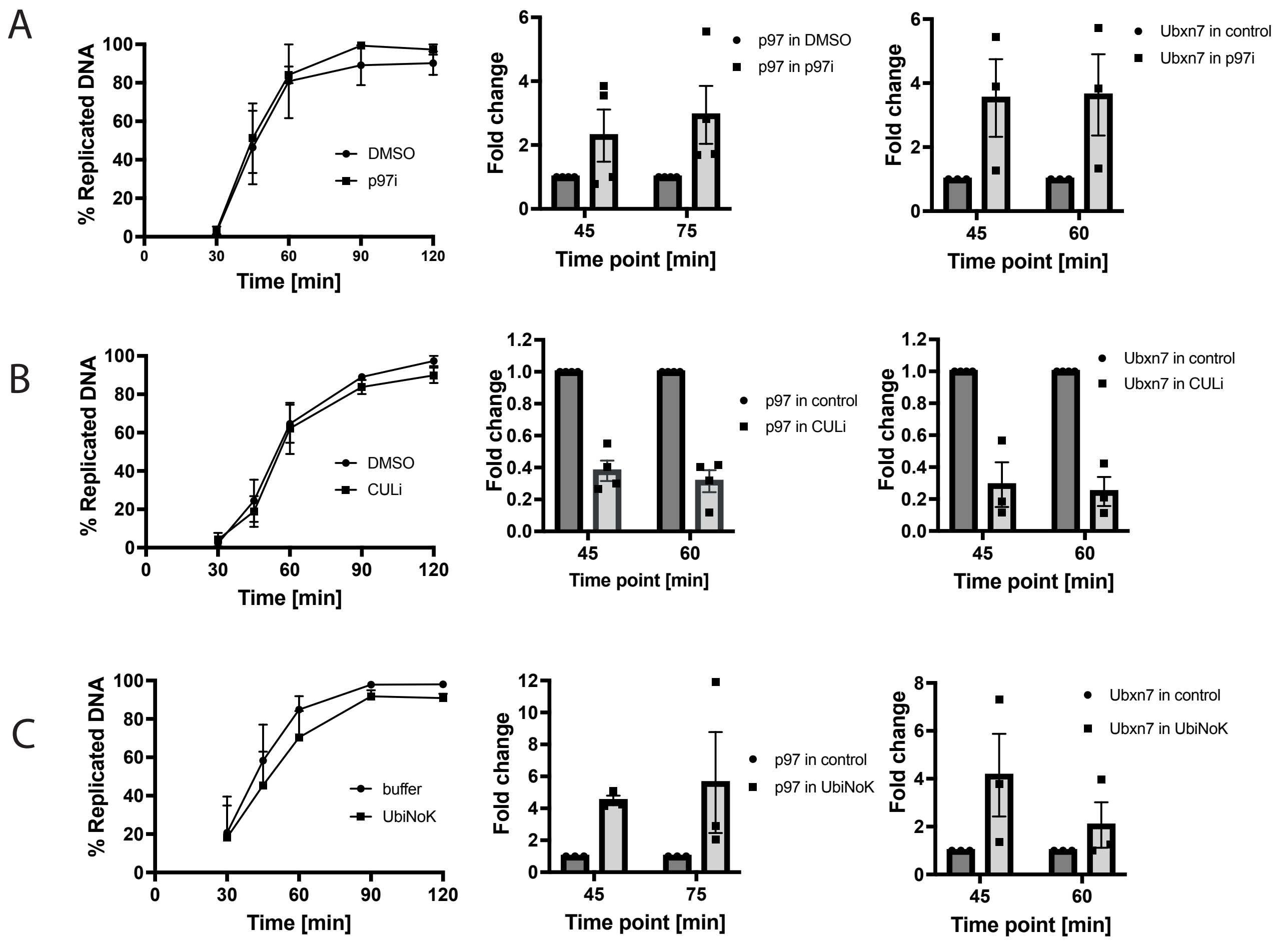


**Supplementary Figure 4**

**(A)** Addition of p97i to egg extract does not affect extract's ability to synthesise DNA. Interphase egg extract was supplemented with DMSO or p97i and the incorporation of $\alpha^{32}$P-dATP into newly synthesised DNA was measured at indicated times. Mean of n=2 for p97i with SEM. p97 and Ubxn7 accumulate on chromatin upon p97i treatment. Quantification of experiment in Figure 4A.

The level of p97 and Ubxn7 bound to chromatin at 45, 60 or 75 min was quantified in DMSO and p97i treated extract. 45 min timepoint represents time when replisomes are present on chromatin in control and treatment sample, while at 60/75 min replisomes are mostly unloaded in control sample. Fold increase in p97i over control is presented as a mean value with individual value points (n=4 for p97 and n=3 for Ubxn7).

**(B)** Addition of CULi to egg extract does not affect extract's ability to synthesise DNA. Interphase egg extract was supplemented with DMSO or CULi and the incorporation of $\alpha^{32}$P-dATP into newly synthesised DNA was measured at indicated times. Mean of n=4 with SEM. p97 and Ubxn7 decreased on chromatin upon CULi treatment - quantification of an experiment in Figure 4B as above (n=4). **(C)** Addition of 6His-UbiNOK to egg extract does not affect extract's ability to synthesise DNA. Analysed as above. Quantification of p97 and Ubxn7 accumulation on chromatin upon 6HIS-UbiNOK treatment as in Figure 4C (n=3).

Tarcan et all Supp Figure 5

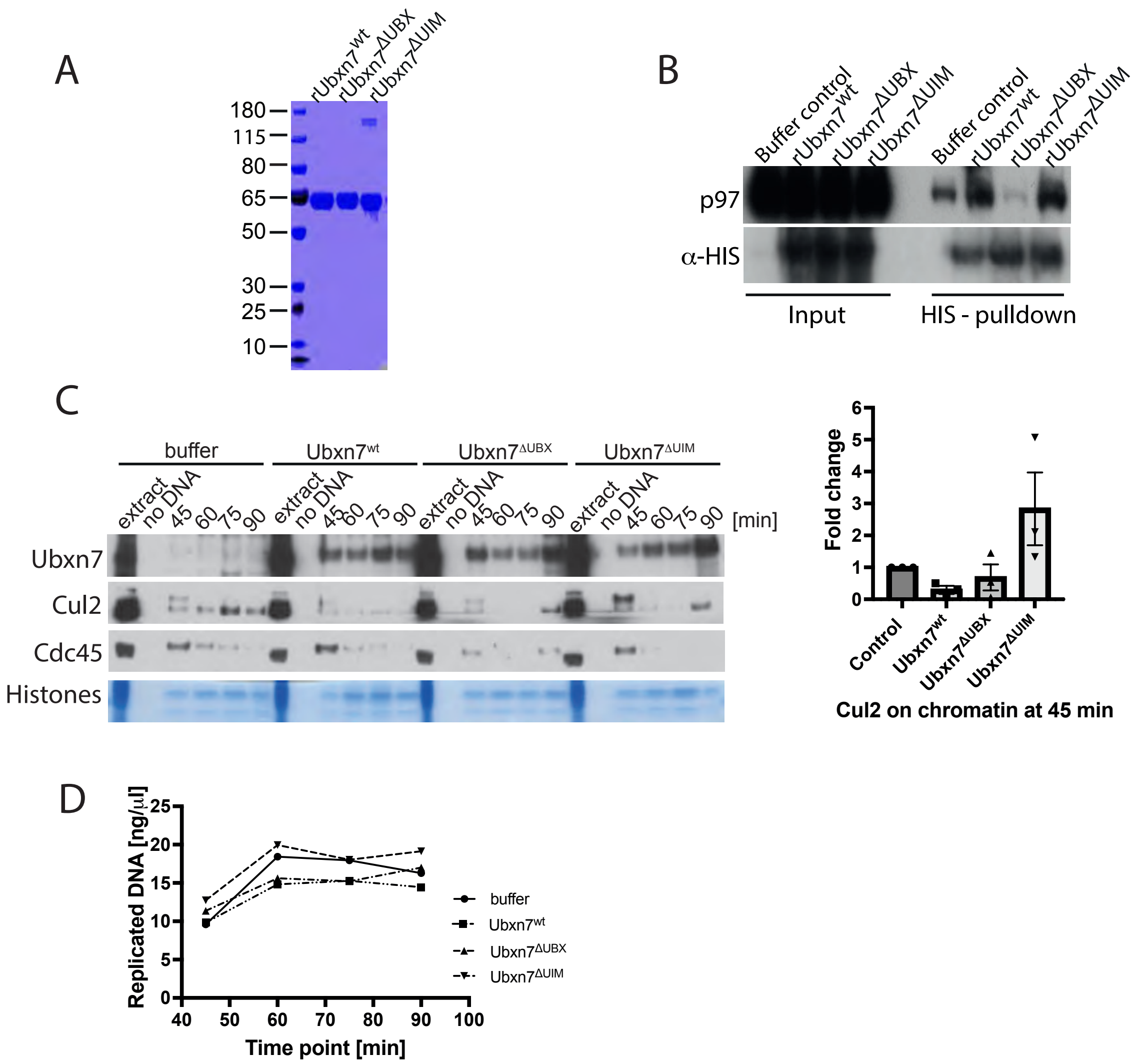


**Supplementary Figure 5.**

**(A)** Recombinant 6xHIS-Ubxn7, 6xHIS-Ubxn7$^{\Delta UBX}$ or 6xHIS-Ubxn7$^{\Delta UIM}$ were purified and equal quantity run on a PAGE gel and stained with coomassie. **(B)** Ubxn7$^{\Delta UBX}$ cannot interact with p97. Interphase egg extract was supplemented with recombinant 6xHIS-Ubxn7, 6xHIS-Ubxn7$^{\Delta UBX}$ or 6xHIS-Ubxn7$^{\Delta UIM}$ and the recombinant proteins were pulled out from replicating egg extract in the middle of S-phase. The ability of recombinant proteins to interact with p97 was analysed by western blotting.

**(C)** Interphase egg extract was supplemented with Ubxn7 or mutants as in (A) and chromatin samples isolated at indicated timepoints during replication reaction. Chromatin samples were analysed by western blotting with indicated antibodies (left). The level of Cul2 bound to chromatin at 45 min of replication reaction was quantified (n=3). Fold change over buffer control is presented. Individual points, mean and SEM. **(C)** Addition of high concentration of rUbxn7 or its mutants to the egg extract does not inhibit egg extract ability to replicate DNA. LFB1/50 buffer or 6HIS-Ubxn7, 6HIS-Ubxn7$^{\Delta UBX}$ or 6HIS-Ubxn7$^{\Delta UIM}$ at 0.3 mg/ml final concenetration. Extract ability to incorporate $\alpha$-$^{32}$PdATP into nascent DNA was quantified.

Tarcan et all Supp Figure 6

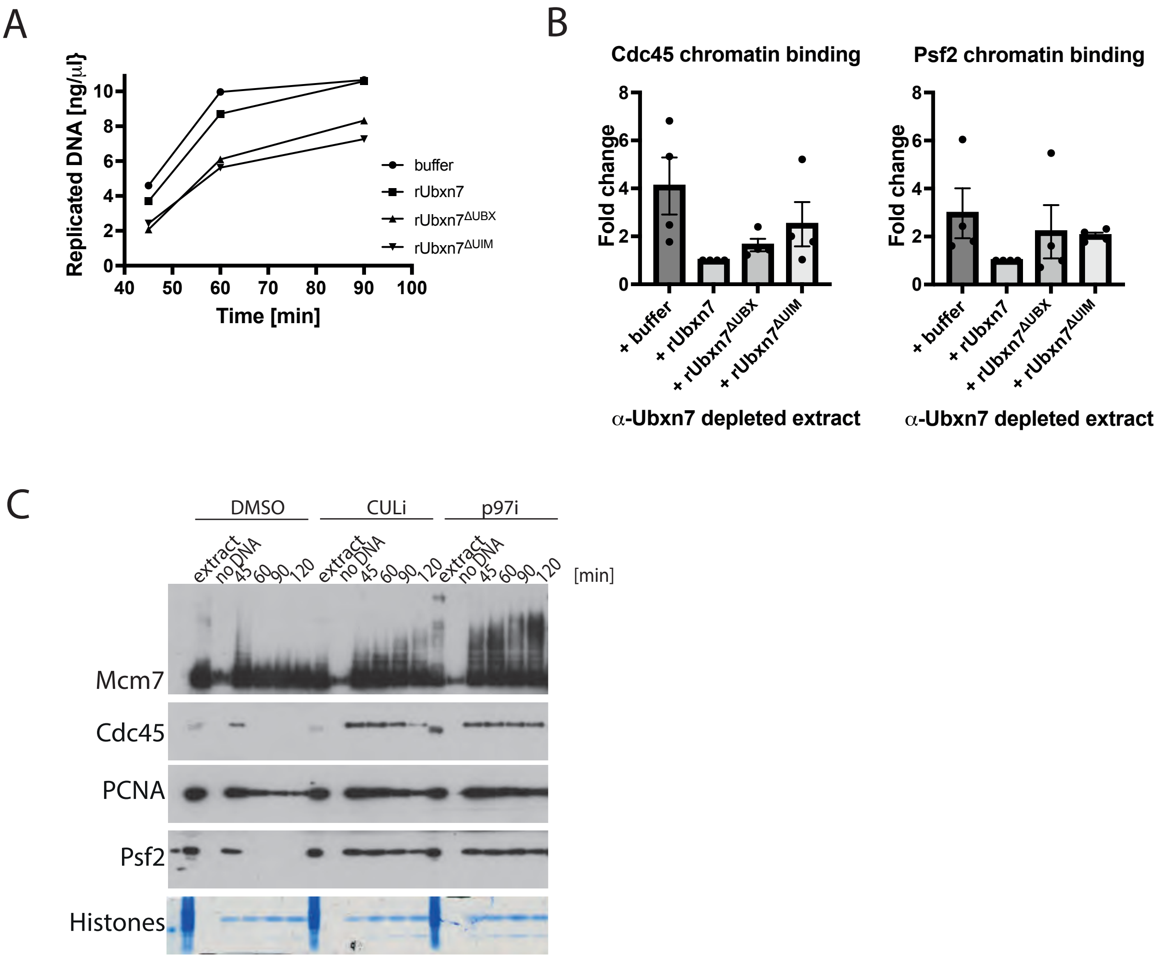


**Supplementary Figure 6.**

**(A)** Addition of recombinant 6xHIS-Ubxn7, 6xHIS-Ubxn7$^{\Delta UBX}$ or 6xHIS-Ubxn7$^{\Delta UIM}$ to rescue Ubxn7-depleted extract does not inhibit extracts ability to synthesise DNA. Ubxn7 depleted extract was supplemented with 6xHIS-Ubxn7, 6xHIS-Ubxn7$^{\Delta UBX}$ or 6xHIS-Ubxn7$^{\Delta UIM}$ and extract ability to incorporate $\alpha$-$^{32}$PdATP into nascent DNA quantified. **(B)** UBX and UIM domains are needed for Ubxn7 activity. Quantification of experiment in Figure 5A. The chromatin bound Cdc45 and Psf2 at 75 min of replication reaction in Ubxn7-depleted extract supplemented with recombinant 6xHIS-Ubxn7, 6xHIS-Ubxn7$^{\Delta UBX}$ or 6xHIS-Ubxn7$^{\Delta UIM}$ were quantified, n=4. Individual points, mean and SEM are presented. **(C)** Mcm7 accumulates on chromatin modified with short ubiquitin chains upon cullin inhibition. Interphase egg extract was supplemented with DMSO, CULi (MLN4924) or p97i (NMS873) and chromatin was isolated during replication reaction at indicated time points after sperm DNA addition. Chromatin samples analysed as in Figure 1A.

Tarcan et all Supp Figure 7

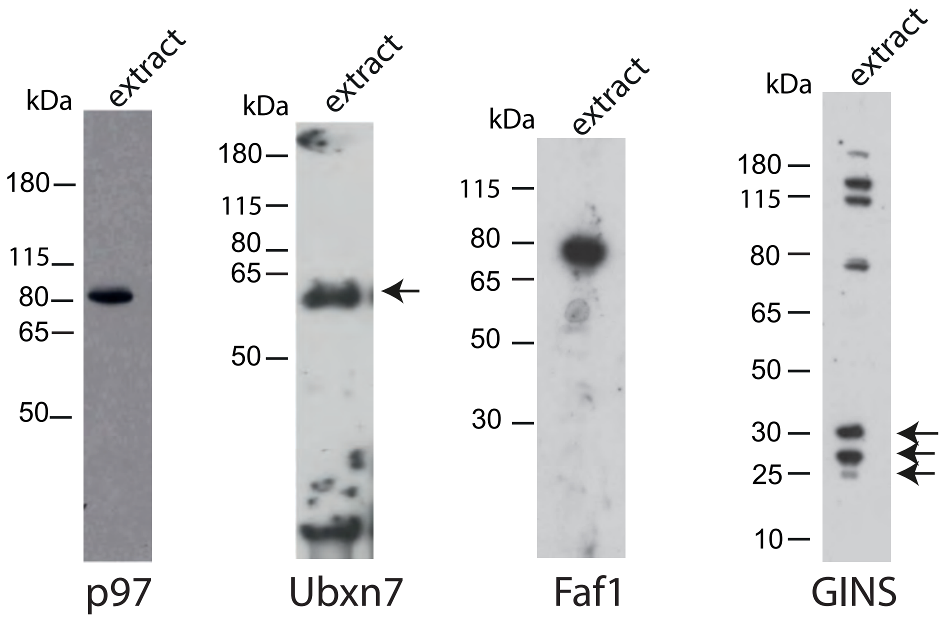


**Supplementary Figure 7**

All of the antibodies raised for this study. For all of the antibodies, a 0.5 µl sample of egg extract was resolved on PAGE and immunblotted using the new affinity purified antibody.

Tarcan et al Table 1

Proteins identified as p97 interactors in experiment described in Figure 1. Factors involved in DNA replication, DNA damage repair and chromatin transactions. Obtained data was compared with mass spectrometry results of interactors of terminated replisomes published previously (4).
Total spectral count is presented with protein coverage in the brackets. Thefull mass spectrometry proteomics data have been deposited to the ProteomeXchange Consortium via the PRIDE repository with the dataset identifier PXD030426 and 10.6019/PXD030426.

| | | IP from chromatin assembled in extract treated with NMS873 | | IP from chromatin assembled in extract treated with p97 mut and caffeine | |
|---|---|---|---|---|---|
| | Protein (kDa) | IgG TSC (coverege) | α-p97 TSC (coverege) | IgG TSC (coverege) | α-Mcm3 TSC (coverege) |
| Mcm Complex | Mcm2 (100 kDa) | 0 | 67 (31%) | 14 (12%) | 957 (72%) |
| | Mcm3 (90 kDa) | 0 | 3 ( 2%) | 22 (10%) | 1133 (84%) |
| | Mcm4 (97 kDa) | 0 | 28 (21%) | 4 (3%) | 945 (78%) |
| | Mcm5 (82 kDa) | 0 | 30 (31%) | 29 (27%) | 862 (84%) |
| | Mcm6 (93 kDa) | 0 | 49 (32%) | 33 (19%) | 828 (83%) |
| | Mcm7 (82 kDa) | 0 | 42 (34%) | 23 (16%) | 884 (77%) |
| Replisome components | Rpa1 (67 kDa) | 0 | 149 (38 %) | 0 | 1 (13%) |
| | Rpa3 (13 kDa) | 0 | 3 (36%) | 0 | 0 |
| | PCNA (29 kDa) | 19 (48%) | 54 (66%) | 0 | 6 (7%) |
| | Dnmt1 (168 kDa) | 0 | 50 (19%) | 0 | 53 (19%) |
| | Spt16 (118 kDa) | 0 | 25 (15%) | 25 (15%) | 0 |
| | Ctf4 (125 kDa) | 0 | 23 (16%) | 0 | 0 |
| | Rfc2 (38 kDa) | 0 | 8 (22%) | 0 | 6 (18%) |
| | Rfc3 (40 kDa) | 0 | 20 (38%) | 4 (11%) | 14 (38%) |
| | Rfc4 (40 kDa) | 0 | 13 (28%) | 0 | 0 |
| | Pol3 (125 kDa) | 0 | 2 (6%) | 0 | 0 |
| | Polα1 (165 kDa) | 0 | 9 (5%) | 0 | 66 (24%) |
| | Polε (261 kDa) | 0 | 3 (1%) | 0 | 316 (37%) |
| | Topbp1-α (169 kDa) | 0 | 11 (71%) | 0 | 0 |
| | Orc2 (62 kDa) | 0 | 8 (12%) | 0 | 39 (29%) |
| | Orc3 (81 kDa) | 0 | 7 (9%) | 0 | 31 (32%) |
| | Orc4 (50 kDa) | 0 | 2 (6%) | 0 | 20 (22%) |
| | Dna2 (120 kDa) | 0 | 6 (5%) | 0 | 0 |
| | Fen1-a (43 kDa) | 0 | 3 (9%) | 0 | 0 |
| DNA replication and DNA damage response proteins | Sall4 (114 kDa) | 0 | 20 (27%) | 0 | 0 (9%) |
| | Rif1 (257 kDa) | 0 | 108 (21%) | 0 | 6 (3%) |
| | Arid1a (206 kDa) | 0 | 82 (17%) | 0 | 0 |
| | Arid1b (148 kDa) | 0 | 16 (10%) | 0 | 0 |
| | Smarca5 (122 kDa) | 0 | 45 (29%) | 2 (2%) | 372 ( 63%) |
| | Smarca4 (181 kDa) | 0 | 21 (9%) | 0 | 0 |
| | Smarce1 (47 kDa) | 0 | 17 (26%) | 0 | 0 |
| | Smarcb1 (43 kDa) | 0 | 11 (24%) | 0 | 0 |
| | Cenpe (339 kDa) | 0 | 41 (6%) | 0 | 0 |
| | Rrm1 (91 kDa) | 0 | 29 (8%) | 0 | 0 |
| | Baz1b (166 kDa) | 0 | 27 (11%) | 0 | 150 (34%) |
| | Rad50 (154 kDa) | 0 | 19 (8%) | 0 | 0 |
| | Rbbp7 (48 kDa) | 0 | 15 (23%) | 0 | 0 |
| | Smc3 (141 kDa) | 0 | 14 (12%) | 0 | 22 (22%) |
| | Hdac1-b (55 kDa) | 0 | 12 (22%) | 0 | 0 |
| | Etaa1 (90 kDa) | 0 | 11 (12%) | 0 | 0 |
| | Ehmt2 (65 kDa) | 0 | 11 (11%) | 0 | 0 |
| | H3.3 (15 kDa) | 4 (21%) | 8 (36%) | 0 | 22 (69%) |
